# Effective Population Size in Field Pea

**DOI:** 10.1101/2024.02.19.581041

**Authors:** Josephine Princy Johnson, Lisa Piche, Hannah Worral, Sikiru Adeniyi Atanda, Clarice J. Coyne, Rebecca McGee, Kevin McPhee, Nonoy Bandillo

## Abstract

**Background:** Effective population size (*N_e_*) is a pivotal parameter in population genetics as it can provide information on the rate of inbreeding and the contemporary status of genetic diversity in breeding populations. The population with smaller *N_e_* can lead to faster inbreeding, with little potential for genetic gain making selections ineffective. The importance of *N_e_* has become increasingly recognized in plant breeding, which can help breeders monitor and enhance the genetic variability or redesign their selection protocols. Here, we present the first *N_e_* estimates based on linkage disequilibrium (LD) in the pea genome.

**Results:** We calculated and compared *N_e_* using SNP markers from North Dakota State University (NDSU) modern breeding lines and United States Department of Agriculture (USDA) diversity panel. The extent of LD was highly variable not only between populations but also among different regions and chromosomes of the genome. Overall, NDSU had a higher and longer-range LD than the USDA that could extend up to 500Kb, with a genome-wide average *r^2^* of 0.57 (vs 0.34), likely due to its lower recombination rates and the selection background. The estimated *N_e_* for the USDA was nearly three-fold higher (*N_e_ =* 174) than NDSU (*N_e_ =* 64), which can be confounded by a high degree of population structure due to the selfing nature of pea.

**Conclusions:** Our results provided insights into the genetic diversity of the germplasm studied, which can guide plant breeders to actively monitor *N_e_* in successive cycles of breeding to sustain viability of the breeding efforts in the long term.

## Introduction

Dry pea (*Pisum sativum* L.), is a diploid, cool-season legume and a member of the Leguminosae family (Abbo et al. 2017). Pea is one of the most important pulse crops grown in more than 100 countries, where 7,043,605 hectares of dry pea were planted around the world with a total production of 12,403,522 tonnes (FAOSTAT 2021; https://www.fao.org/faostat/). In the USA alone, the pea production reached one million tonnes in 2019 (USDA 2020). In recent years, pea protein has become more popular in the market for plant-based diets e.g., Beyond® Meat Burger (Bari et al. 2021). Pea seeds have earned a reputation as a dietary goldmine with around 15 – 32% protein content, vitamins, folate, fibers, potassium and minerals, which is good for human health and helps prevent cardiovascular and specific cancer diseases (Bari et al. 2021; Tayeh et al. 2015). The increasing popularity of plant-based proteins in the market has further propelled the demand for peas. Therefore, the study of genetic diversity should expand to accelerate the genetic gain of pea varieties to meet future demands, maintaining the diversity in peas is the top priority for plant breeders (Bari et al. 2021; Gali et al. 2019).

Estimation of effective population size (*N_e_*) determines the rate of inbreeding (Rahimmadar et al. 2021; Tenesa et al. 2007) and genetic changes due to genetic drift (Gargiulo et al. 2023). *N_e_* is an important parameter in population genetics and breeding introduced by Sewall Wright in 1931, which helps breeders to maintain and monitor the level of genetic diversity in their species (Cobb et al. 2019). The estimated *N_e_* is expected to be smaller than the census size (*N*), as it influences the rate at which genetic diversity decreases within a population (Lonsinger et al. 2018; Hare et al. 2011). Relatively smaller *N_e_* indicates limited population diversity, which, in turn, can restrict genetic advancement within a breeding program (Hayes et al., 2003). Moreover, *N_e_* parameter retrieves the population dynamics of the genes (Nei and Tajima 1981).

The effective size of a population refers to the hypothetical number of individuals in an idealized population that would exhibit a comparable genetic response to stochastic processes, similar to that observed in a real-world population which is based on the Wright-Fisher model (Wang et al. 2016; Wright 1931; Fisher 1930). This model shows genetic drift as the main operating factor, and that changes in allelic and genotypic frequencies over generations are solely influenced by the population size (*N*) (Wang et al. 2016). In real-world breeding populations, factors such as mutation, migration, natural selection, and non-random mating come into play (Wang et al. 2016) These factors affect the actual rates of inbreeding and changes in gene frequency variance observed in a population (Charlesworth 2009). This will indeed impact *N_e_* and therefore, reduce the genetic variation and diversity. The most commonly used extensions for effective population size theory are variance effective size and inbreeding effective size (Wang et al. 2016). The variance effective size reflects the rate of change in gene frequency variance, while inbreeding effective size corresponds to the rate of inbreeding observed in a population (Crow and Kimura 1970). These measures allow us to quantify the consequences of genetic drift in a real population, based on the characteristics and dynamics of the idealized Wright-Fisher population (Wang et al. 2016).

While *N_e_* of a population can be estimated either from demographic data or genetic markers, the latter is preferred (Gilbert and Whitlock 2015; Luikart et al. 2010; Fernández et al. 2005). Demographic data involves using census size and variance of reproductive success whereas genetic markers reveal changes in allele frequencies over time and are based on linkage disequilibrium (LD). When the pedigree or demographic data is not available, *N_e_* can be estimated using genetic markers (Wang 2005). The most popular and widely-employed genetic approach has been the temporal method, which relies on temporal fluctuations in allele frequencies observed on multiple samples collected from the same population (Nei and Tajima 1981). *N_e_*, however, can also be directly estimated using LD between loci at various distances along the genome (Hayes et al. 2003; Sved 1971). Recent advancements in high-throughput sequencing and the availability of high-density markers such as single nucleotide polymorphisms (SNPs) have increased over the past decade, contributing to the LD-based approach now being acknowledged as more reliable, robust (Novo et al. 2022), cost and time effective than the temporal approach (Gargiulo et al. 2023).

Linkage disequilibrium (represented as *r^2^*) is a phenomenon characterized by the non-random association of alleles at various loci (Hill and Robertson 1968) which became popular in recent years for predicting *N_e_* (Antao et al. 2011). Correlations between alleles are generated by genetic drift when it is inversely proportional to *N_e_* (Gargiulo et al. 2023), which changes the allele frequencies in a population over time. The biggest advantage of LD over the temporal method (Pollak 1983), is the strength of associations between markers that can be used to calculate *N_e_* at any time (generations) from a single population accurately without relying on longitudinal data. This makes LD a valuable tool for studying populations where temporal information may be limited or unavailable. Recombination and mutation rates are fundamental processes that shape the genetic landscape (Ardlie et al. 2002), and by analyzing LD, we can better understand their history and apply it to plant breeding and population genetics (Sved and Hill 2018).

In this study, we estimated the extent of LD decay in the dry pea genome and utilized the relationship between LD and recombination frequency, as initially described by Sved (1971), to estimate *N_e_* which is convenient as it only requires one sampling time (García-Cortés et al. 2019; Hill 1981). We used two sets of populations: 1) NDSU modern breeding lines, hereafter referred to as NDSU set, and 2) USDA diversity panel, hereafter referred to as USDA set. Our objectives were two-fold: (i) to estimate *N_e_* for these two germplasms set in dry pea and (ii) to compare the genetic variation between these germplasms. To achieve these goals, we developed a comprehensive R package that implements the Sved (1971) formula for *N_e_* prediction. This package not only caters to the specific needs of dry pea research but can also be adapted for use in other crop species. Since there has been no information on *N_e_* for peas, our findings serve as a valuable reference for researchers seeking to determine the minimum number of lines required for designing experiments. Furthermore, comparing the genetic variation between NDSU modern breeding lines and USDA multi-environmental lines provides valuable information about the diversity and potential of these germplasm collections. This knowledge can guide breeding programs and conservation efforts, ensuring the maintenance and enhancement of genetic resources in dry pea cultivation.

## Methods

### Plant Materials

In this study, we used plant materials from two distinct germplasms. The first population comes from the NDSU Pulse Breeding Program (NDSU set) where 300 advanced elite lines were generated from multiple bi-parental populations. These lines were created specifically with a focus on phenotypes including high yield, grain quality, resistance to disease and some other desirable agronomic traits. The breeding lines used in this experiment were carefully chosen and contain both contemporary and past elite germplasm. (Bari et al. 2023; Atanda et al. 2022).

The second population is from a USDA diversity panel (USDA set), and contained 482 accessions, of which 292 samples were from the Pea Single Plant Plus Collection (Pea PSP) (Bari et al. 2021; Holdsworth et al. 2017; Cheng et al. 2015). The USDA set was composed of accessions that represent most of available diversity within the USDA pea germplasm collection based on the knowledge of geography, taxonomy, morphology and genotyping-by-sequencing data generated previously (Holdsworth et al. 2017).

### DNA extraction, Sequencing and Variant Calling

Leaf tissues from the greenhouse were collected at different stages for all NDSU elite lines and USDA accessions. The DNA from the lyophilized tissues were extracted using the DNeasy Plant Mini Kit (Qiagen). Detailed information regarding the tissue collections and extractions are provided in Bari et al. (2023) and Bari et al. (2021). Both NDSU set and USDA set were sequenced using genotyping-by-sequencing (GBS). Using the restriction enzyme *ApeKI*, dual-indexed GBS libraries for both populations were prepared (Elshire et al. 2011). Samples were sequenced using NovaSeq S1 × 100 Illumina sequencing technologies. The NDSU set sequenced libraries were retrieved with a quality score ≥ 30. For USDA set, FASTQC (Andrews 2010) was utilized to perform quality check and removed reads with lengths < 50 bases. All reads that passed the quality check were aligned with the reference genome (Kreplak et al. 2019) (https://www.pulsedb.org). Finally, the aligned reads were analyzed using SAMtools (v1.10) and generated the variant files (VCF) using FreeBayes (V1.3.2).

The amount of single nucleotide polymorphisms (SNPs) identified for the NDSU set was 28,832, while 380,527 SNP markers were identified in the USDA set (Bari et al. 2021, 2023; Atanda et al. 2022). For these marker datasets, we filtered minor allele frequency (MAF), since alleles with < 5% could produce bias to the LD and *N_e_* calculations (Toosi et al. 2010, Lee et al. 2014). We also removed markers with more than 20% missing values using Plink v1.9 (Purcell et al. 2007) and heterozygosity > 20% using Tassel v5.0 (Bradbury et al. 2007). The resulting marker sets consisted of 7,157 (NDSU set) and 19,826 (USDA set) SNP markers that were used for downstream analysis.

### Calculation of Linkage Disequilibrium (*r*^2^)

LD was calculated using Plink v1.9 (Purcell et al. 2007) with a maximum distance of 750 kb. Using “ggplot2” R package, the genome-wide and chromosome-wide LD-decay (*r*²) were visualized against the physical distance (kb) to show the recombination history (see Fig. 1 & 2).

**Figure 1.**
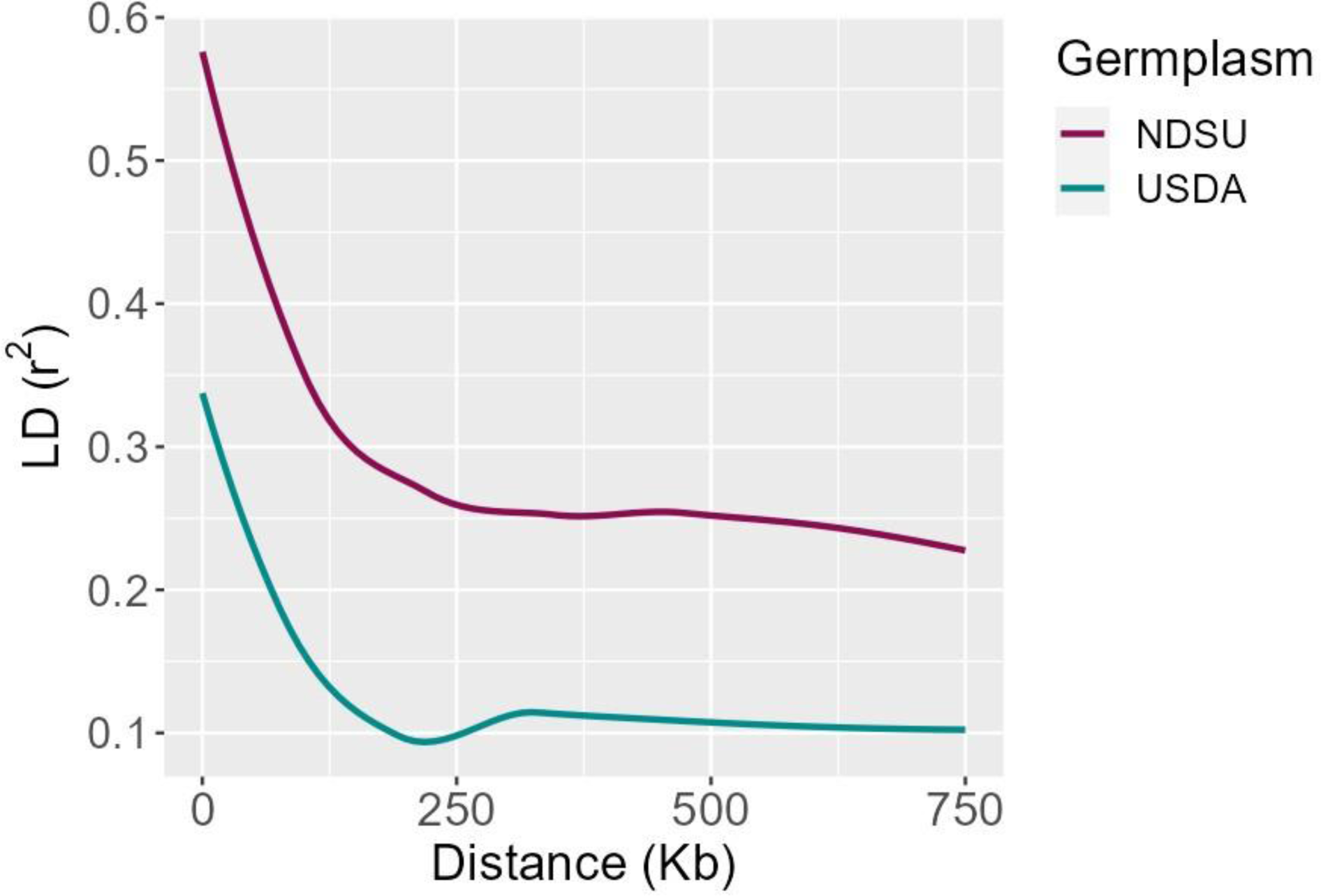
Genome-wide Linkage Disequilibrium - decay of NDSU set and USDA set

LD scores were also estimated using Genome-wide Complex Trait Analysis (GCTA) software for window size of 1000 kb and *r*^2^ cutoff of 0 (Yang et al. 2011). This approach was employed to visualize the distribution of mean LD throughout the genome.

### Calculation of Effective Population Size

Effective population size (*N_e_*) for both the NDSU set and the USDA set were estimated based on LD using the Sved (1971) equation. The recombination rate (cM) was calculated using cM/Mb conversion ratio from a recent pea genetic linkage map (Sawada et. al. 2022) and then transformed to Morgan’s (*c*).

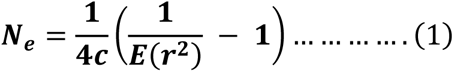

Where, *N*_*e*_ = effective population size

*c* = genetic distance in Morgan’s

*E*(*r*^2^) = expected *r*^2^

The expected *r^2^*was predicted by linear regression model using least square estimation (LSE), Prediction of *r*^2^:

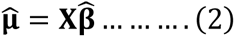

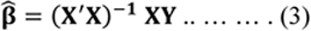

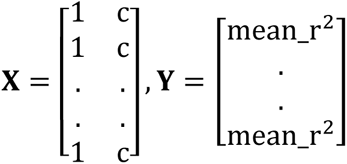

The mean_*r*^2^ from the ***Y*** parameter was calculated by LD (*r^2^*) for the genetic distance ‘c*’* using ‘group by’ mean function in R Environment (R Core Team, 2023).

Now with the availability of all required parameters, we finally estimated *N_e_* from Equation (1) using LSE,

According to the formula (Eqn. 1), we assigned the variables as predictor (**X**) and response (**Y**) and calculated the coefficient **β**_1_ without the intercept term **β**_0_, following Juma et al. (2021).

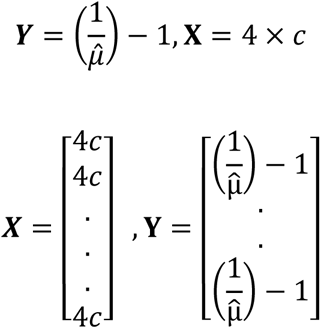

Again, we used Equation (3) to calculate the coefficient **β**_1_ which represents *N_e_*.

## Results

### Linkage Disequilibrium Decay Rate and Scores

The decay of linkage disequilibrium (*r^2^*) was examined in both NDSU set and USDA set by utilizing 7,157 and 19,826 SNP markers, respectively. This analysis allowed for the identification of the physical distance at which the decay rate occurred. Supplementary Figure 1 depicts the distribution of SNPs within and across chromosomes for both populations, providing an illustration of the marker density. The NDSU set’s genome-wide LD-decay plot (Figure 1) demonstrates that the *r^2^* reached its peak value of 0.57 within the initial kilobases and subsequently exhibited a gradual decline. The *r^2^* showed a decrease from 0.3 to 0.25 when the genomic distance increased from 150 kb to 250 kb. Following that, the LD within each chromosome was observed visually in Figure 2 in order to improve comprehension of the decay pattern. Chromosomes 1 and 6 exhibited a rapid decay at approximately 175 kb, while chromosomes 2 and 5 demonstrated a comparatively slower decay rate of around 350 kb. Furthermore, it is worth noting that chromosome 5 had the higher *r^2^* value of 0.61 compared to other chromosomes. Whereas, the genome-wide LD of USDA set showed that *r^2^* started at a lower value of 0.34 and dropped rapidly and reached 0.2 and 0.1 at 100 kb and 200 kb (Figure 1). From the chromosome-wide LD-decay (Figure 2), we observed that chromosome 3 dropped faster around ∼150 kb, but the *r^2^* decreased below 0.1 for chromosomes 4 and 7. Also, chromosomes 1, 5 and 6 decayed slowly (∼250 kb) and reached *r^2^* 0.1. We also observed that chromosome 1 exhibited a higher *r^2^* of 0.37. LD-decay figures show the trend of the *r^2^*decaying from LD to linkage equilibrium (LE).

**Figure 2.**
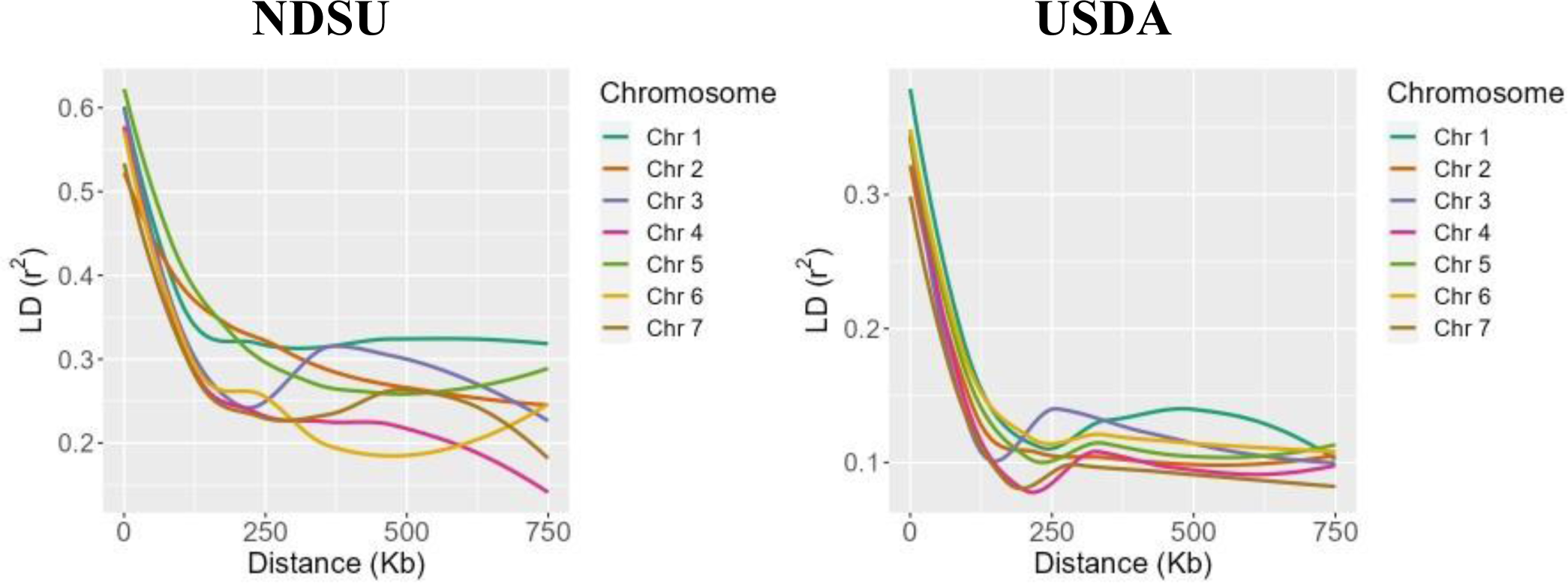
Chromosome-wide Linkage Disequilibrium - decay of NDSU set and USDA set

Additionally, we performed calculations of LD scores as an alternative metric for inferring LD. The analysis of local LD in the NDSU set indicates a notable rise in the average *r^2^* of 0.6 across all chromosomes. The average *r^2^* of chromosomes 5 and 6 was the highest with 0.8. The genomic interval encompassing the centromeric region of chromosome 2 was missing. In contrast, the USDA set exhibited low average *r^2^*, with chromosome 2 hardly reaching 0.4, and chromosomes 1, 4, and 7 having few sets that reached 0.3. It is worth noting that the LD density of the NDSU set is comparatively lower than the USDA set (Figure 3).

**Figure 3.**
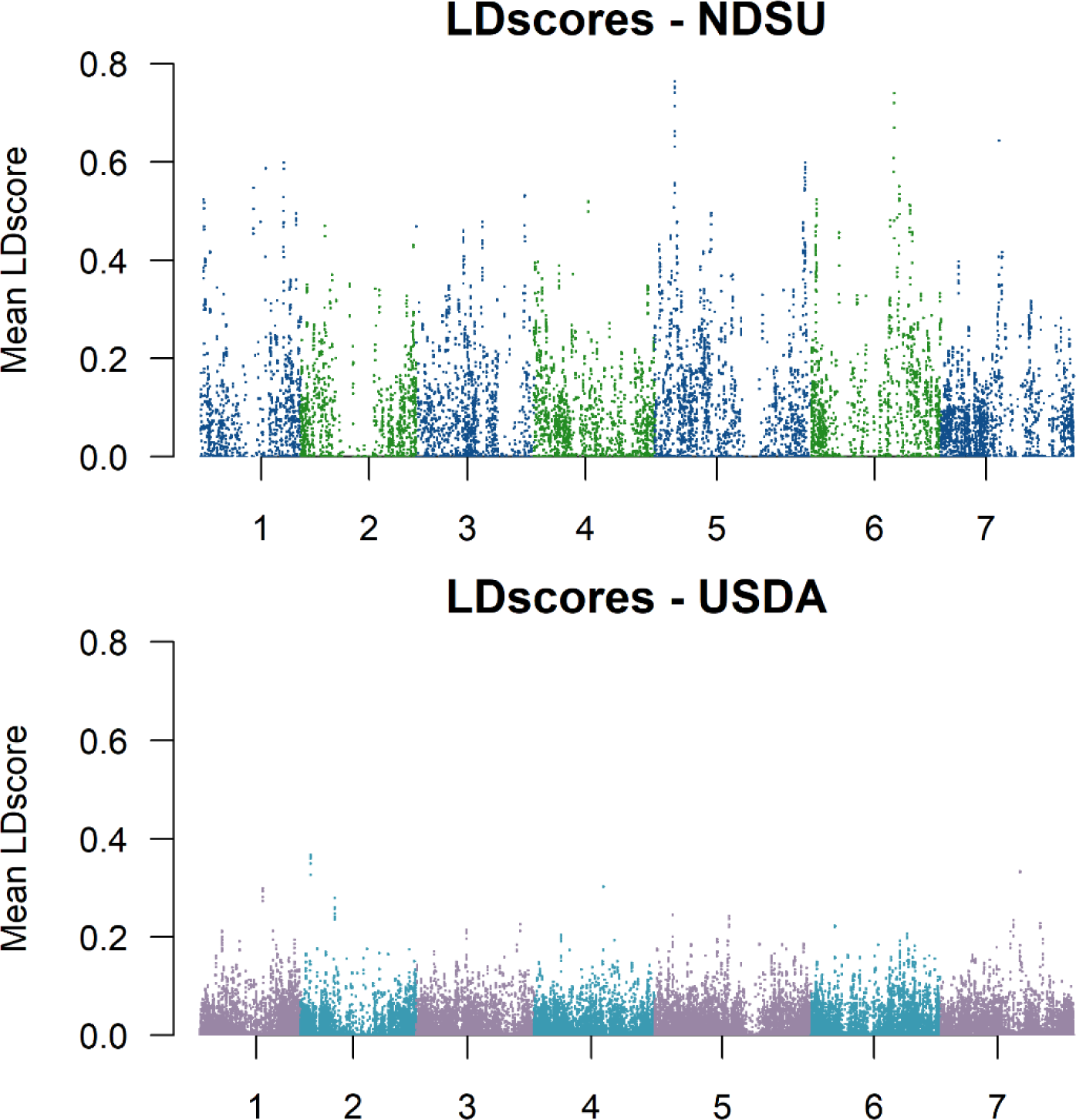
The Mean LD scores estimated in 1000kb windows. There is a significant increase in LD of NDSU set compared to USDA set

With respect to recombination rate (centimorgans - cM), the genome-wide *r^2^* on average decayed from 0.54 to 0.27 at 0.7 cM for the NDSU set, indicating a moderate level of correlation within this specific genetic distance across the genome. In contrast, the USDA set had lower average *r^2^*(0.28) which dropped within a shorter genetic distance (0.5 cM). This implies that as the distance between the markers increases to 0.5 cM, they tend to be less correlated with each other (Supplementary Figure 2)

The level of LD exhibited significant variation across distinct genomic regions and populations of dry peas. The impracticality of conducting whole-genome scanning can be attributed to the excessive number of markers required for such studies, particularly in cases where there is a low level of linkage disequilibrium (Kruglyak 1999). The USDA set reported a low LD value, indicating a higher occurrence of recombination events. In contrast, the NDSU set showed a higher LD score, suggesting a greater frequency of linked markers presumably due to limited recent recombination to date (Siol et al. 2017).

### Effective Population Size (*N_e_)*

Based on LD, the estimated effective population size (*N_e_*) for both the populations are shown in Figure 4. The smaller *N_e_* and high LD in NDSU set indicates that it has undergone selective pressures leading to reduced diversity and increased correlation between the markers. Given NDSU set’s population history and marker density, it is acceptable to state that despite lower *N_e_*, it holds a reasonable level of diversity that may help maintain its genetic variability which is essential for long-term viability and adaptability. The USDA set resulted in lower LD and higher *N_e_*, meaning it has more diversity and has encountered relatively fewer instances of selective pressures or genetic bottlenecks. It is important to note that the low LD can also be observed in a population with high *N_e_*. Thus, it was expected to see NDSU set with lower *N_e_* compared to USDA set. These estimates explain how genetic drift and selections have shaped these populations over time.

**Figure 4.**
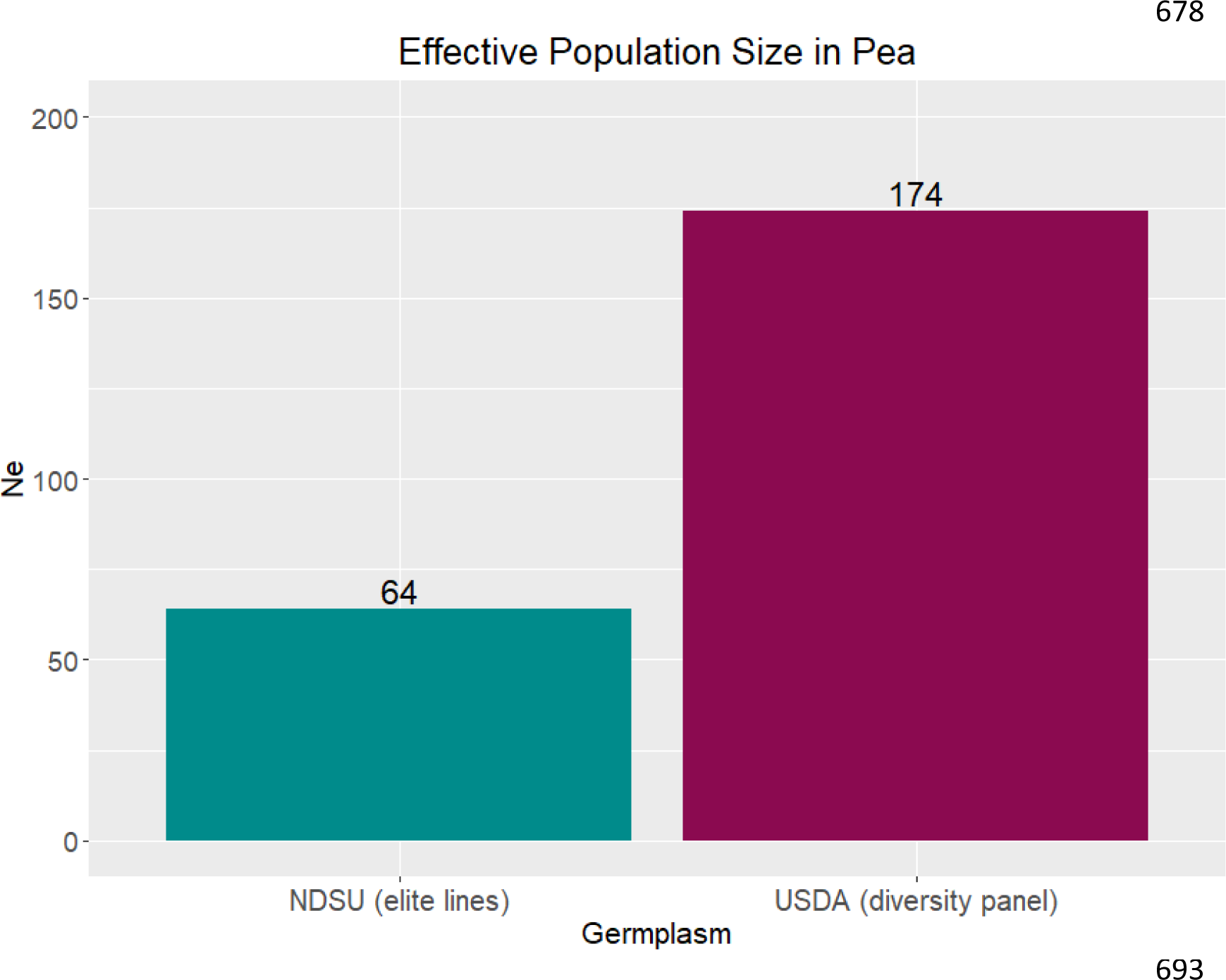
Estimated effective population size (*N_e_*) for NDSU set is 64 and USDA set is 174

## Discussion

The importance of *N_e_* has become increasingly recognized in plant breeding as it describes the rate of inbreeding and can reflect the contemporary status of genetic diversity in breeding populations (Onda and Mochida 2016). When *N_e_* is low, the population can become quickly inbred with little potential for genetic gain making long-term selection ineffective. Therefore, plant breeders should be cognizant of the effective population size of their breeding program (Cobb et al. 2019). Actively monitoring *N_e_* in successive cycles of breeding can enhance the viability of the breeding efforts and help sustain long-term genetic gain. In this study, we presented the first estimation of *N_e_* in dry pea using two distinct germplasm sets: 1) the NDSU set consisting of elite breeding lines within the NDSU breeding program, and 2) the USDA set comprised of landraces and plant introductions collected all over the world (Cheng et al. 2015; Holdsworth et al. 2017). The former represents breeding lines and germplasm in an active breeding program that releases new modern cultivars, while the latter represents germplasm accessions in a repository. As expected, the estimated *N_e_* for the USDA set (*Ne*=174) was higher than the NDSU set (*Ne*=64). The genetic diversity for the USDA set is higher than the NDSU set as it represents most of the available diversity in the USDA pea germplasm collection (Holdsworth et al. 2017; Cheng et al. 2015).

The *N_e_* estimate for the NDSU set was within the same range as those reported in other self-pollinating crops such as rice (*Oryza sativa*) and soybean (*Glycine max*), with calculated *N_e_* ranging from 20 to 60. Juma et. al. (2021) estimated the *N_e_* in rice to be 22 using an elite core panel comprised of 72 lines, but *N_e_* may have been underestimated due to limited marker information used in the analysis. Similar studies in rice also had the same range of *N_e_*, with calculated values ranging from 23-57 and 40-60; these were estimated based on breeding populations from recurrent selection programs (Grenier et al. 2015) and pedigree data (Morais Júnior et al. 2017). The estimated *N_e_* of USDA set was within the range of *N_e_* values reported in studies conducted on other crops. In soybean, Xavier et al. (2018) estimated *N_e_* for the USDA soybean germplasm collection comprised of 19,652 accessions from Bandillo et al. (2015) and reported it to be 106 individuals. Recent studies have shown that soybean possess several genetic bottlenecks (Guo et al. 2010) and its genetic diversity has been reduced (Li et al. 2013, Min et al. 2010). The *N_e_* estimate of USDA set is relatively higher than soybean, implying greater diversity. Zhao et al. (2013) estimated *N_e_* in wild rice using 11 Chinese *Oryza rufipogon* populations including 32 landraces and reported it between 96-158, which is in a similar range to the USDA set. Thus, the *N_e_* of USDA set offers greater potential for adaptation, maintaining rare alleles, population stability, and reduced risk for inbreeding.

The results of our study also suggest that the use of GBS holds good potential for making inferences of *N_e_* regardless of the germplasm type. Using GBS-based markers, we approximated the LD pattern within and across chromosomes of both germplasms and then used the LD information for estimation of *N_e_*. Genome-wide LD (*r^2^*) of the USDA set decayed from lower LD at 200 kb, while the NDSU set had the highest LD declined at a longer distance of around 250 kb. These results provided consistency of higher genetic variations of the former over the latter. Similar LD findings have been observed in previous studies conducted on peas, wherein both wild and spring peas exhibited a decay distance of approximately 200 kb, whereas wild/landrace peas were around 100 kb (Siol et al. 2017) which is a bit lower than the USDA set. Comparing the LD of USDA set and the NDSU set to other selfing crops such as rice, soybeans, and barley, the physical distances found were more or less similar depending on the populations. For instance, Huang et al. (2010) estimated LD using *O. indica* and *O. japonica* landraces of rice at 123 and 167 kb, respectively, with *r^2^* declining to 0.25 and 0.28. Additionally, soybean landraces extended from 90 to 500 kb (Hyten et al. 2007) while improved cultivars hit 133 kb (Zhou et al. 2015) which is similar to the USDA set. Alternatively, a recent LD analysis from soybean USDA germplasm revealed that the *r^2^* dropped intragenically within a few kilobases (Xavier et al. 2018) and the one in barley’s landraces hit 90 kb (Caldwell et al. 2006), both shorter than the USDA set. The LD-decay of the NDSU set was also found to be in a similar range with elite varieties of barley which extended to at least 212 kb (Caldwell et al. 2006) and *O. japonica* elite lines at ∼318 kb (Li et al. 2020), but had a higher distance compared to *O. indica* elite lines (∼124 kb) (Li et al. 2020). The LD-decay rate of a crop does depend on the genetic background of the populations being studied, and it can be affected due to mutations, genetic drift, non-random mating, and a small *N_e_* (Flint-Garcia et al. 2003).

Since public plant breeding programs are moving toward more quantitative methods, the importance of the dynamic exchange of genetic material and the maintenance of diversity within the population has increased. Effective population size helps breeders preserve and remodel their selection strategies to enhance the stability and variability in their breeding populations (Cobb et al. 2019). Breeders can also implement marker-based mating experiments known as optimum contribution selection (OCS) (Juma et al. 2021) in order to maintain diversity in selection candidates for long-term gain. As pulse crop breeders navigate through challenges in their breeding programs, the information from this study provides valuable insights by demonstrating the strength of contemporary populations and possibly contributing to the long-term goal of increasing genetic gain while maintaining diversity in breeding programs.

## Conclusions

These research findings shed light on the range of genetic diversity in both NDSU set and USDA set. The evaluation of *N_e_* can be a bit more challenging and there is a possibility of potential biases if certain crucial factors including sample size, marker density, population history and LD are not accounted appropriately (Waples and Yokota 2007, Waples and Do 2010; Gilbert and Whitlock 2015; Marandel et al. 2020). Even though genetic markers have become a more widely utilized approach for estimating *N_e_* in recent years, there are still more obstacles to overcome in its *N_e_* accuracy. Future estimation of *N_e_* could be complemented with gene expression along with DNA markers and demographic history, that would increase the understanding of breeders regarding the population dynamics and potential for adaptation in different environments.

## Acknowledgments

The authors would like to acknowledge the funding provided by USDA-NIFA (Hatch Project #: ND01513). The genotyping of the NDSU materials was funded by the North Dakota Department of Agriculture through the Specialty Crop Block Grant Program (19-429) and Northern Pulse Growers Associations. The genotyping of the USDA germplasm was partially supported through funding from USDA Plant Genetic Resource Evaluation, USA Dry Pea and Lentil Council Research Committee, USDA ARS Pulse Crop Health Initiative and USDA ARS Project: 5348-21000-017-00D (CJC), and 5348-21000-024-00D (RJM). This investigation used resources of the Center for Computationally Assisted Science and Technology (CCAST) at North Dakota State University, Fargo, ND, USA which were made possible in part by NSF MRI Award No. 2019077.

We would also like to acknowledge the contributions of Jérôme Bartholomé who provided technical guidance on the implementation of Sved’s (1971) equation.

## Author Contributions

**Josephine Princy Johnson:** Conceptualization; Data curation; Pipeline development; Formal analysis; Investigation; Methodology; Writing – original draft, review and editing, **Lisa Piche:** Methodology; Review and editing, **Hannah Worral:** Methodology; Review and editing, **Sikiru Adeniyi Atanda:** Writing - review and editing, **Clarice J. Coyne:** Funding acquisition; Resources; Review and editing, **Kevin McPhee:** Funding acquisition; Resources; Review and editing, **Rebecca McGee:** Funding acquisition; Resources; Review and editing, and **Nonoy Bandillo:** Conceptualization; Supervision; Funding acquisition; Resources; Validation; Writing - review and editing.

## Data archiving

Please find the “EffectivePopSize” *R* package from GitHub repository: https://github.com/PrincyJohnson/EffectivePopSize.

