## Supplementary Figures 1 & 2 for "Effective Population Size in Field Pea"

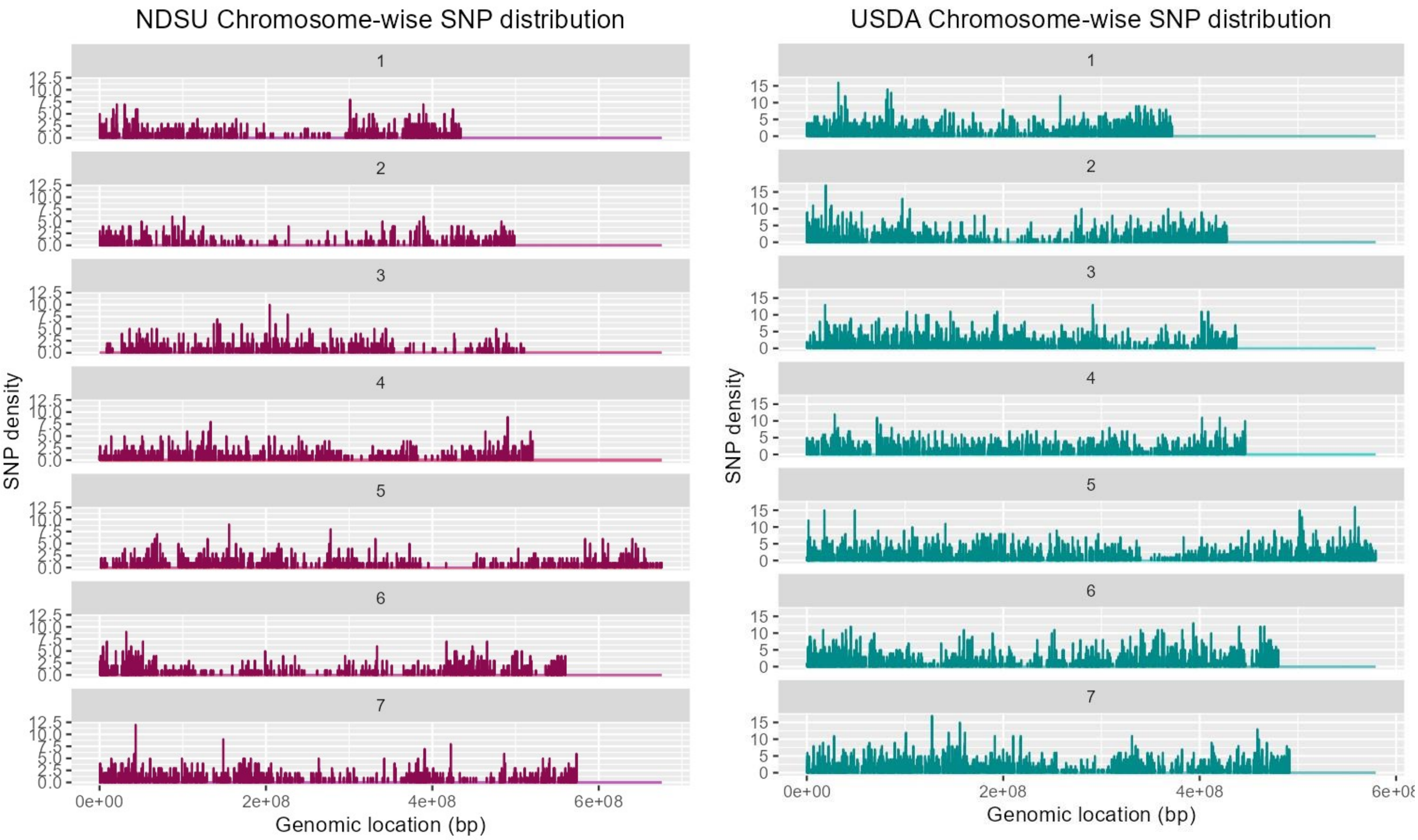


Supplementary Figure 1: SNP density of NDSU set and USDA set, x-axis is the genomic location (bp) and y-axis is the density

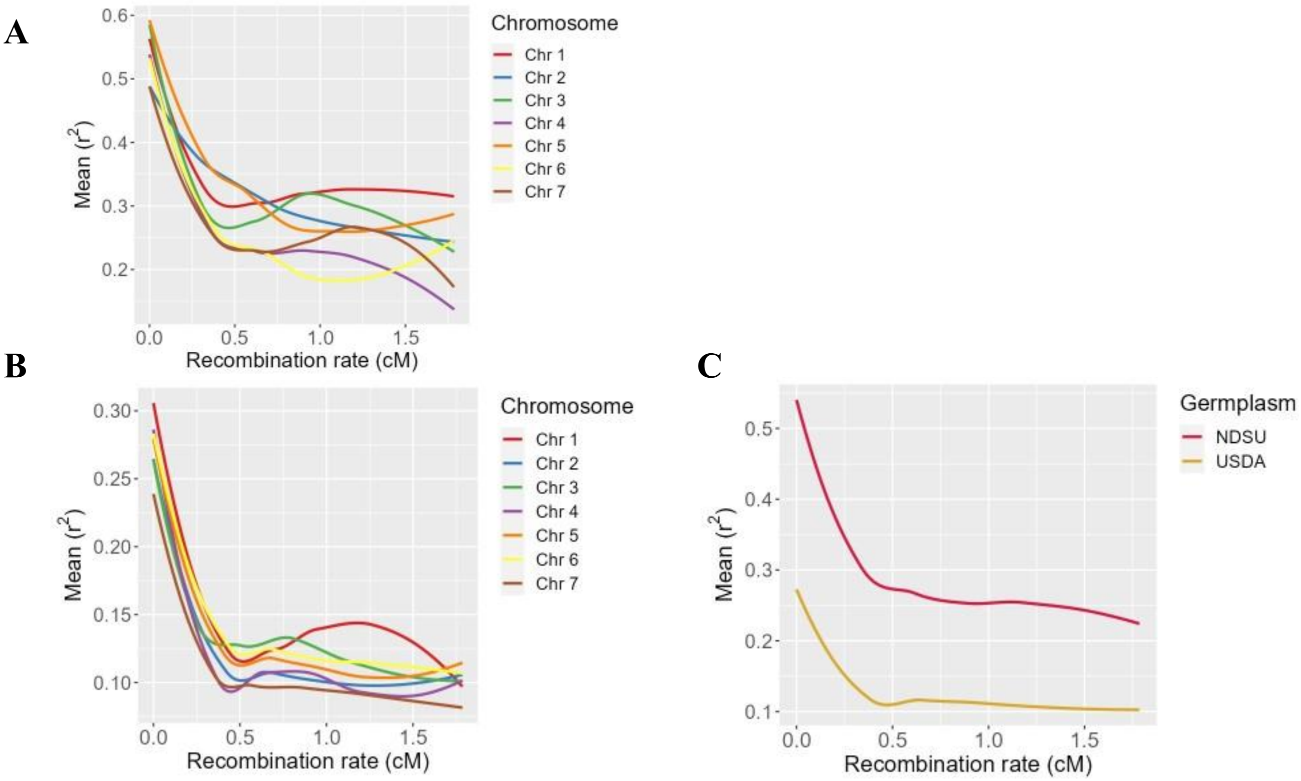


Supplementary Figure 2: Genome-wide (C) and Chromosome-wide linkage disequilibrium decay in the NDSU set (A) and USDA set (B) with mean of $r^2$ (y-axis) and recombination rate (cM) (x-axis)
